# Improving Thermal and Gastric Stability of Phytase via pH Shifting and Coacervation: A Demonstration of Bayesian Optimization for Rapid Process Tuning

**DOI:** 10.1101/2025.04.18.649602

**Authors:** Waritsara Khongkomolsakul, Poompol Buathong, Eunhye Yang, Younas Dadmohammadi, Yufeng Zhou, Peilong Li, Lixin Yang, Peter I. Frazier, Alireza Abbaspourrad

## Abstract

Phytase (phyA) breaks down phytate, which can help with nutrient absorption in a plant-based seed diet or high-phytate food. Unfortunately, it is prone to denaturation at food preparation temperatures and is easily inactivated by pepsin during gastric digestion. To protect phyA for use in high-temperature processes (100 °C) and gastric digestion, chitosan (CS) was used to complex phyA. Bayesian optimization, a machine learning technique, was used to demonstrate how to expedite the optimization process. Thermal stability of the optimized complex increased from 20% (Control: phyA in the native state) up to 74% at 4:1 CS to phyA (CS-phyA) complex and 52% at the 1:1 CS-phyA complex as measured by phytase activity assay. Chitosan complexation also improved the retention of enzyme activity after thermal and gastric digestion by 13-fold, retaining residual activity at 40% for the 4:1 CS to phyA and 22% for the 1:1 CS-phyA complexes compared to the enzyme itself, which only retained 3% residual activity. Molecular docking and circular dichroism were used to investigate the underlying interaction mechanism between CS and phyA and the secondary structure of the enzyme after heat treatment. Confocal laser scanning microscopy (CLSM) and scanning electron microscopy (SEM) confirmed the complexation of phyA with CS and revealed complex morphology. With improved enzyme stability, there is great potential for efficiently expanding phytase applications in a high plant-based seed food matrix.

**Graphical Abstract:** 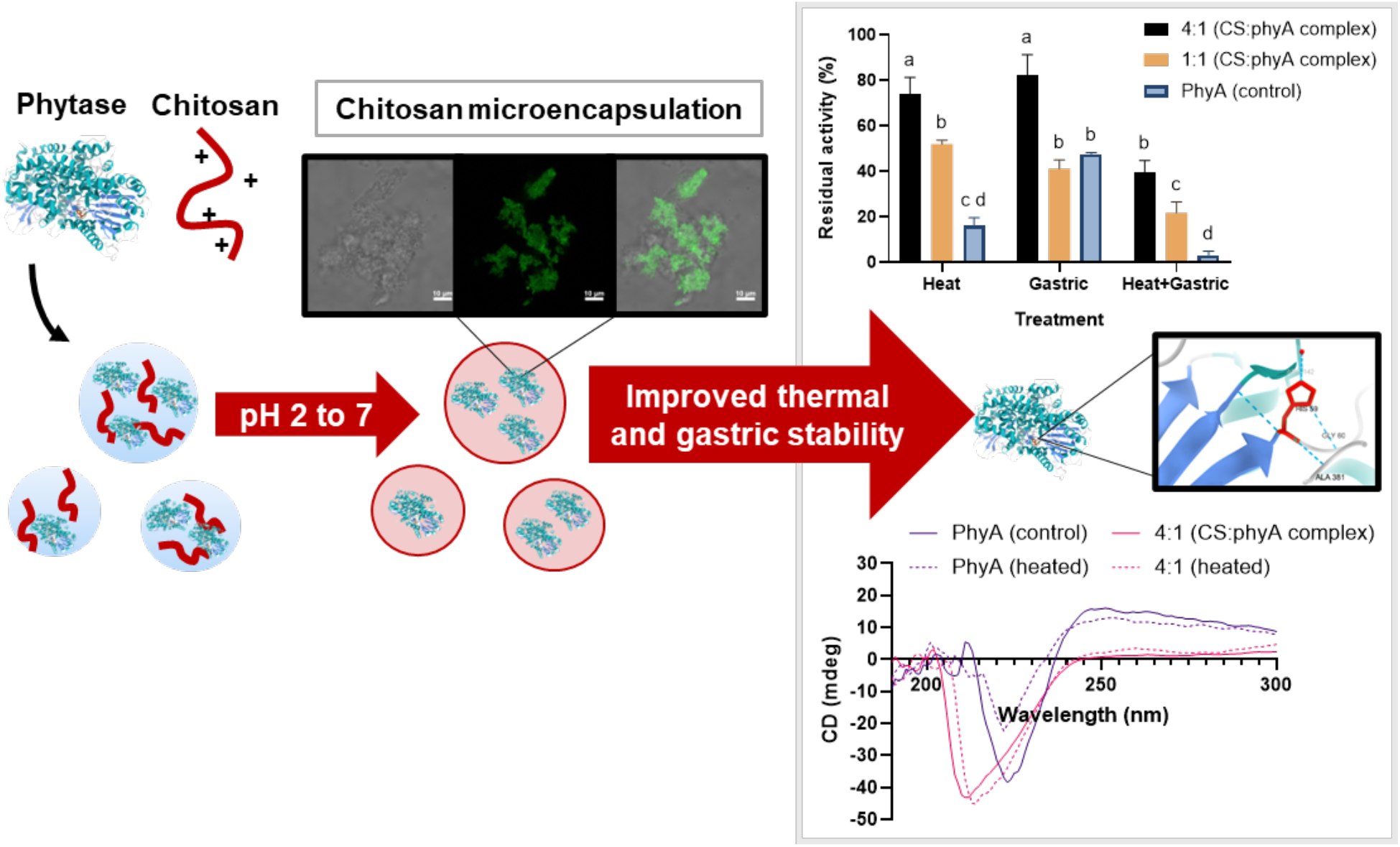

**Highlights:**

- Thermal stability of phytase improved from 20% to 74% using chitosan
- pH shifting increased enzyme complexation efficiency and thermal stability (100 °C)
- Bayesian optimization (BO) is a promising optimization tool for complexation conditions

## 1. Introduction

Enzymes play a crucial role in enhancing the nutritional value of food by increasing the absorption of micronutrients. Phytase (myo-inositol-hexakisphosphate, phyA) has gained attention for its ability to break down phytic acid, a micronutrient inhibitor, thus improving the bioavailability of essential nutrients such as iron, zinc, and calcium (Cowieson & Ravindran, 2007). The human body has phyA in only small amounts, which are insufficient to break down the amount of phytic acid present in plant-based foods, thus, micronutrient absorption is inhibited in the upper part of the gastrointestinal tract (Iqbal et al., 1994). The ability of dietary phytase has been assessed as a pathway to increase micronutrient absorption. Numerous clinical trials have aimed to determine the effectiveness of phytase supplementation in enhancing micronutrient absorption, specifically iron (Gupta et al., 2015; Troesch et al., 2009). It was proven that integrating phytase with high-phytate food, such as maize porridge, nearly doubled the iron absorption in women. However, difficulty arises in integrating phytase into the food matrix as phytase activity is influenced by external stressors, such as high temperature or acidic conditions, and consequently decreases phytase’s ability to improve the absorption of micronutrients (Monnard et al., 2017).

Enzyme microencapsulation, complexation, and immobilization are widely used in the food industry and pharmaceuticals to enhance enzyme stability for food processing and drug delivery. One promising strategy is engineering an electrostatic interaction between the enzyme and biopolymer (Lin et al., 2022). Electrostatic interaction is often used to form a coacervate complex through the optimization of pH level and enzyme to biopolymer ratio. Therefore, the pH level and its consequent impact on the chosen target enzyme and biopolymer must be considered.

Chitosan (CS), is a natural non-toxic biopolymer that can be harvested from the fungi *Aspergillus niger* as an alternative to animal sources (Abdel-Gawad et al., 2017). The structure of CS is composed of D-glucosamine (GlcN) and N-acetyl-D-glucosamine (GlcNAc) units crosslinked with beta-1,4 glycosidic bonds, and depending on the percent deacetylation, the ratio between GlcN and GlcNAc may vary. Chitosan has free amino and hydroxyl groups that influence its overall charge, making its pKa about ~6-7. Thus, it is often dissolved in acidic conditions and then used with negative polysaccharides or proteins during complexation (Hosseinnejad & Jafari, 2016). Several studies have reported an improved resistance to thermal degradation using CS on other enzymes, such as lipase (Q. Li et al., 2024; Rafiee & Rezaee, 2021). A study by Liu et al. showed improved lipase activity across different pH levels and at elevated temperatures of 70 °C (Liu et al., 2023). This elevated enzyme thermal stability afforded by complexation, however, may not be sufficient to protect against environmental stress involved in food processing, such as cooking, where temperatures around 100 °C for at least 10 minutes are often required.

Coacervation has been proven to enhance protein stability, but the optimization conditions can influence its protective effect on the enzyme or target protein (Lin et al., 2023). A promising approach to improve solubility and emulsifying properties is pH shifting, which directly affects the hydrophobicity, disulfide bonds, and surface charges of the protein or biopolymer (Dong et al., 2024). Therefore, integrating and optimizing pH shifting with coacervate formation could maximize protection against thermal degradation. To find the optimal pH and the appropriate ratio of CS and phyA for the best protection against thermal degradation and to maintain the activity of phyA, it is essential to sample the optimization combination carefully. To maximize efficiency in the optimization process and to provide a blueprint for future work, using or developing appropriate statistical methods is essential.

Bayesian optimization (BO) (Frazier, 2018; Mockus, 1975) has emerged as a versatile framework for optimizing expensive-to-evaluate and black-box objective functions. BO has demonstrated its efficiency across a wide range of applications, including hyperparameter optimization for machine learning models (Snoek et al., 2012), protein engineering (Hu et al., 2023), pharmaceutical product development (Sano et al., 2020), and materials design (Frazier & Wang, 2016), among others. The primary goal of BO is to identify the global maximum with the fewest possible experiments. In brief, BO uses seed data (D_n_) to build a surrogate model using a Gaussian process (GP) to predict the system’s trend. BO then used this fitted model to construct an acquisition function, which is used to evaluate the benefits of performing an additional experiment under any possible conditions. BO then suggested the most promising condition that maximizes the benefits. An experiment with this suggested condition is conducted, and the new data is subsequently added to the dataset. The process repeats until we reach the global maximum or the target result.

We investigated the impact of pH shifting on the formation of electrostatic complexes between phyA and CS and evaluated their stability, thermal stress, and gastric degradation. We evaluated the impact of pH changes and CS to phyA ratio on complexation efficiency, yield, and loading capacity using the bicinchoninic acid protein assay (BCA). We then tested the enzyme activity after thermal treatment at 100 °C for 12 min and under typical gastric conditions (Yang et al., 2024). We used BO to optimize experimental factors *x*, specifically pH and ratio, for CS complexation to find the optimal combination with the highest thermal stability *f*(*x*). Circular dichroism and molecular docking were used to confirm and further understand the complex mechanism, and SEM and CLSM were used to explore the complex morphology. As this initial study shows, the use of Bayesian optimization can be generalized and used for future coacervate complex optimization to accelerate research in food.

## 2. Materials and methods

### 2.1 Materials

Phytase from *Aspergillus niger* (phyA) was provided by DSM company (Heerlen, Netherlands). Chitosan from *Aspergillus niger* (MW 300kDa, >98% deacetylation) was purchased from Sarchem Laboratories (Farmingdale, NJ, US). Ammonium molybdate tetrahydrate, acetic acid glacial, Fluorescein 5(6)-isothiocyanate (FITC), L-ascorbic acid, Pepsin lyophilized salt-free from porcine (~2500 units/mg protein), trichloroacetic acid (TCA), and sodium acetate were purchased from Sigma-Aldrich (St. Louis, MO, USA). BCA protein assay kit (the pierce™ rapid gold), hydrochloric acid, and sodium hydroxide were purchased from Thermo Fisher Scientific (Rockford, IL, USA). Sulfuric acid (H_2_SO_4,_ >95%) was obtained from Honeywell Fluka (Charlotte, NC, USA). Sephadex G-25 was purchased from Cytiva (Marlborough, Massachusetts).

### 2.2 Chitosan-phyA complex formation and optimization

#### 2.2.1 Preparation of complex

Solutions of phytase (phyA) (0.3 w/v%) were prepared by diluting the stock solution provided by the DSM company with Milli-Q water. Chitosan was dissolved in acetic acid (pH 5.0) at room temperature (~25 °C). Both phyA and CS solutions were individually adjusted to the starting pH levels between 2.0 and 7.0 with either 1 M NaOH or 1 M HCl. After pH adjustment, CS and phyA solutions were mixed at different ratios (CS:phyA): 1:12, 1:8, 1:6, 1:4, 1:2, 1:1, 2:1, 4:1, 6:1, 8:1, and 12:1. The solutions were then mixed on a rotator at 24 rpm for 30 min and adjusted to the final pH levels between 5.0 and 7.0 to form CS-phyA complexes. The solutions were centrifuged at 10,000*g* for 10 min at 4 °C. The supernatant was used for protein quantification to calculate complexation efficiency, and the pellet, which contains the CS-phyA complex, was collected and stored at –20 °C for at least 12 hours. The frozen complex was then lyophilized overnight or until completely dry. The final complex was collected in a powder form for further characterization analysis.

#### 2.2.2 Bayesian optimization for pH-ratio combination for forming the complex

A Bayesian optimization workflow was created to recommend the next CS:phyA ratio at which to perform experiments based on any provided dataset toward the goal of maximizing thermal stability. In this workflow, the dataset is 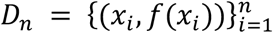, where *n* is the number of datapoints. Each input *x*_*i*_ represents the ratio of CS to phyA in a complexation system, and *f*(*x*_*i*_), denotes our objective function: the average measured thermal stability over three replicates. This experiment considers 11 discrete CS:phyA ratios: 1:12, 1:8, 1:6, 1:4, 1:2, 1:1, 2:1, 4:1, 6:1, 8:1, and 12:1. These discrete ratios are encoded into a continuous real number by the amount of CS per unit phyA, forming the search space of ratios *χ* = {0.08, 0.13, 0.17, 0.25, 0.50, 1.00, 2.00, 4.00, 6.00, 8.00,12.00}. The objective, thermal stability, was measured using the activity assay described. We chose to optimize based on thermal stability, as the purpose of complexation is to improve phyA thermal stability. Our BO workflow uses a Gaussian process (GP) model (Rasmussen & Williams, 2006). This GP uses a constant prior mean *μ*_0_(*x*) = *C* and the Matérn-5/2 covariance function:

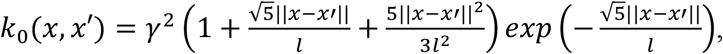

where *γ* and *l* are kernel hyperparameters. These kernel hyperparameters, *γ* and *l*, and the constant mean, *C*, are estimated by the Maximum Likelihood Estimation (MLE) method. The prior mean function represents the initial expectation of the objective function values across the input space. The covariance function, also called the kernel, encodes assumptions about the structure of the function, such as smoothness, and models the correlation between function values *f*(*x*) and *f*(*x*′) at any pair of inputs *x* and *x*′ in the search domain.

The dataset *D*_*n*_ is incorporated into the GP model using Bayes’ theorem, resulting in a posterior distribution over the function values. The posterior mean *μ*_*n*_(*x*|*D*_*n*_), which provides the predicted objective function value at any input *x*, is given by

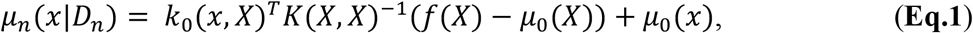

The posterior variance 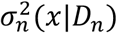, which quantifies the uncertainty of the prediction, is given by

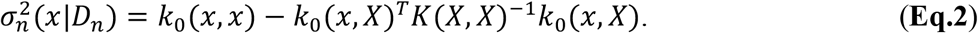

Here *X* = [*x*_1_ *x*_2_ … *x*_*n*_] is the vector of *n* inputs that have been evaluated, *k*_0_(*x, X*)^*T*^ = [*k*_0_(*x, x*_1_) *k*_0_(*x, x*_2_) … *k*_0_(*x, x*_*n*_)],

*K*(*X, X*) is the covariance matrix with elements *K*(*X, X*)_*i,j*_ = *k*_0_(*x*_*i*_, *x*_*j*_),

*μ*_0_(*X*)^*T*^ = [*μ*_0_(*x*_1_) *μ*_0_(*x*_2_) … *μ*_0_(*x*_*n*_)] and

*f*(*X*)^*T*^ = [*f*(*x*_1_) *f*(*x*_2_) … *f*(*x*_*n*_)].

To illustrate the use and behavior of GP models, a GP fit is used to predict thermal stability value at a fixed pH of 2.0 across varying CS:phyA ratios (**Figure 1**). The model is trained using observations obtained at the 11 distinct ratios, represented by red dots in the figure. The posterior mean function *μ*_*n*_(*x*|*D*_*n*_) is depicted as the solid blue line, while the associated 95 % confidence interval, defined as *μ*_*n*_(*x*|*D*_*n*_) ±1.96× *σ*_*n*_(*x*|*D*_*n*_), is shown as a shaded blue region.

**Figure 1.**
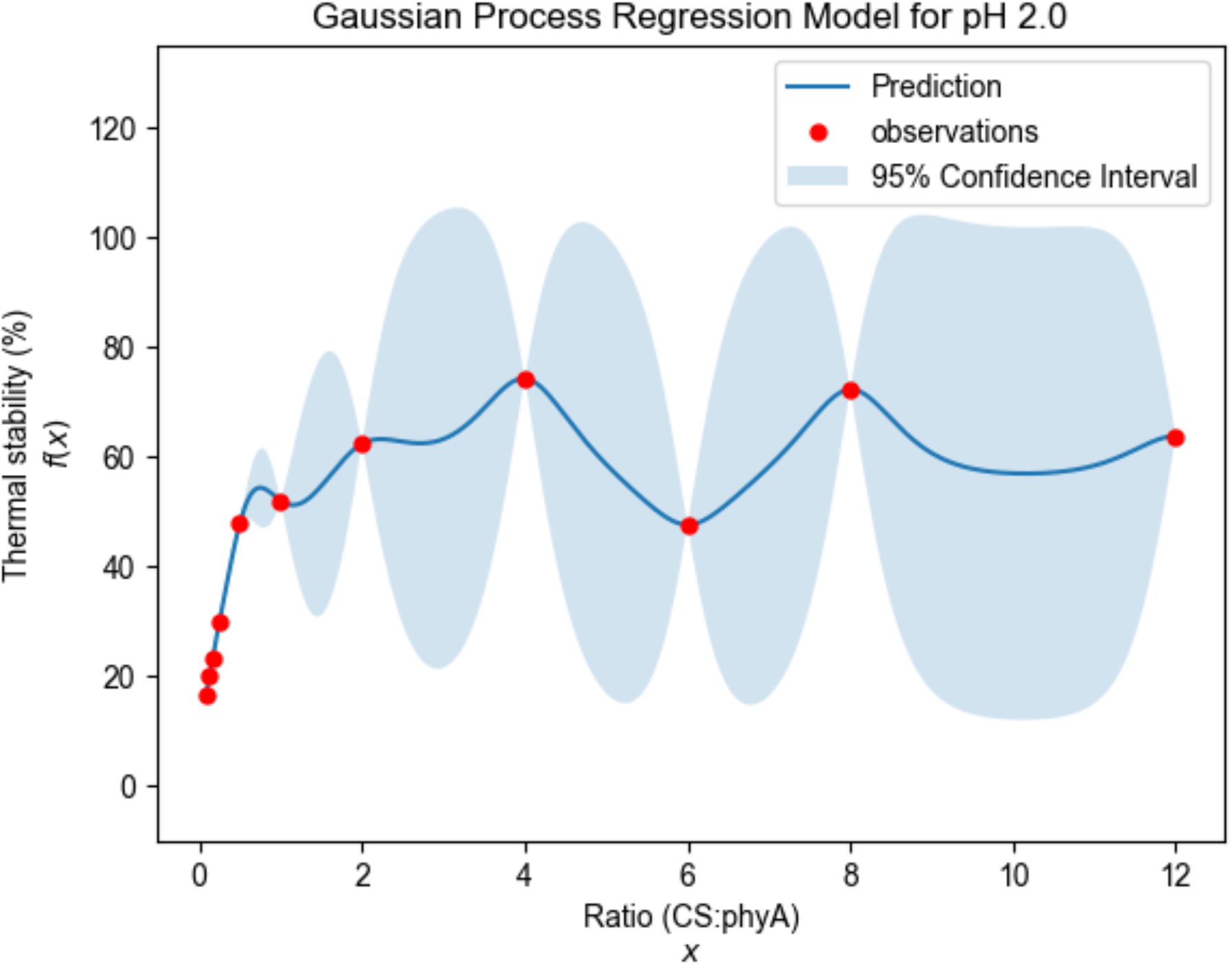
A Gaussian process regression model was fitted to a specific thermal stability dataset at pH 2.0.

In the created BO workflow, after fitting the GP model, BO then uses an acquisition function to evaluate the benefit of conducting an additional experiment. For our acquisition function, we use the Expected Improvement (EI) (Jones et al., 1998)

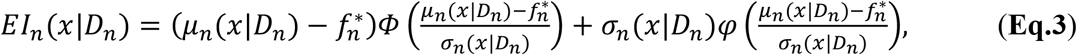

where 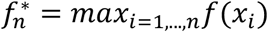, is the best current objective function value found so far, *Φ* and *φ* are the cumulative distribution function (CDF) and probability density function (PDF), respectively, of a standard normal random variable, and *μ*_*n*_(*x*|*D*_*n*_) and *σ*_*n*_(*x*|*D*_*n*_) are the predictive value and uncertainty, respectively, from **Eqs. 1** and **2**.

In the workflow, based on the GP fit, BO suggests a CS:phyA ratio *x*^∗^ in the search space domain *χ*, that maximizes the EI acquisition function to conduct an experiment:

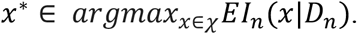

Intuitively, maximizing the Expected Improvement (EI) acquisition function balances the trade-off between exploitation and exploration strategies based on the posterior distribution of the fitted GP model. Exploitation refers to selecting input values in regions where the posterior mean value *μ*_*n*_(*x*|*D*_*n*_) is high, typically near input that has previously yielded the best observed objective value 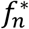. This results in a local search around the current best input. Exploration, in contrast, promotes sampling in regions where the posterior uncertainty *σ*_*n*_(*x*|*D*_*n*_) is large, typically areas that have no previously evaluated inputs, since potentially better solutions may exist in areas that have not been thoroughly evaluated.

To implement the BO workflow created as described above, an experiment would be performed at the suggested ratio input *x*^∗^, and we would obtain the associated thermal stability value *f*(*x*^∗^). This new data pair (*x*^∗^, *f*(*x*^∗^)), would be subsequently added to the dataset, and the process would repeat until we reach the target requirement, such as 50% thermal stability improvement on phyA (>70% residual activity).

**Figure 2** summarizes the BO workflow, emphasizing its iterative nature. We implement the algorithm in this workflow using the BoTorch package in Python (https://botorch.org/) (Balandat et al., 2020).

**Figure 2.**
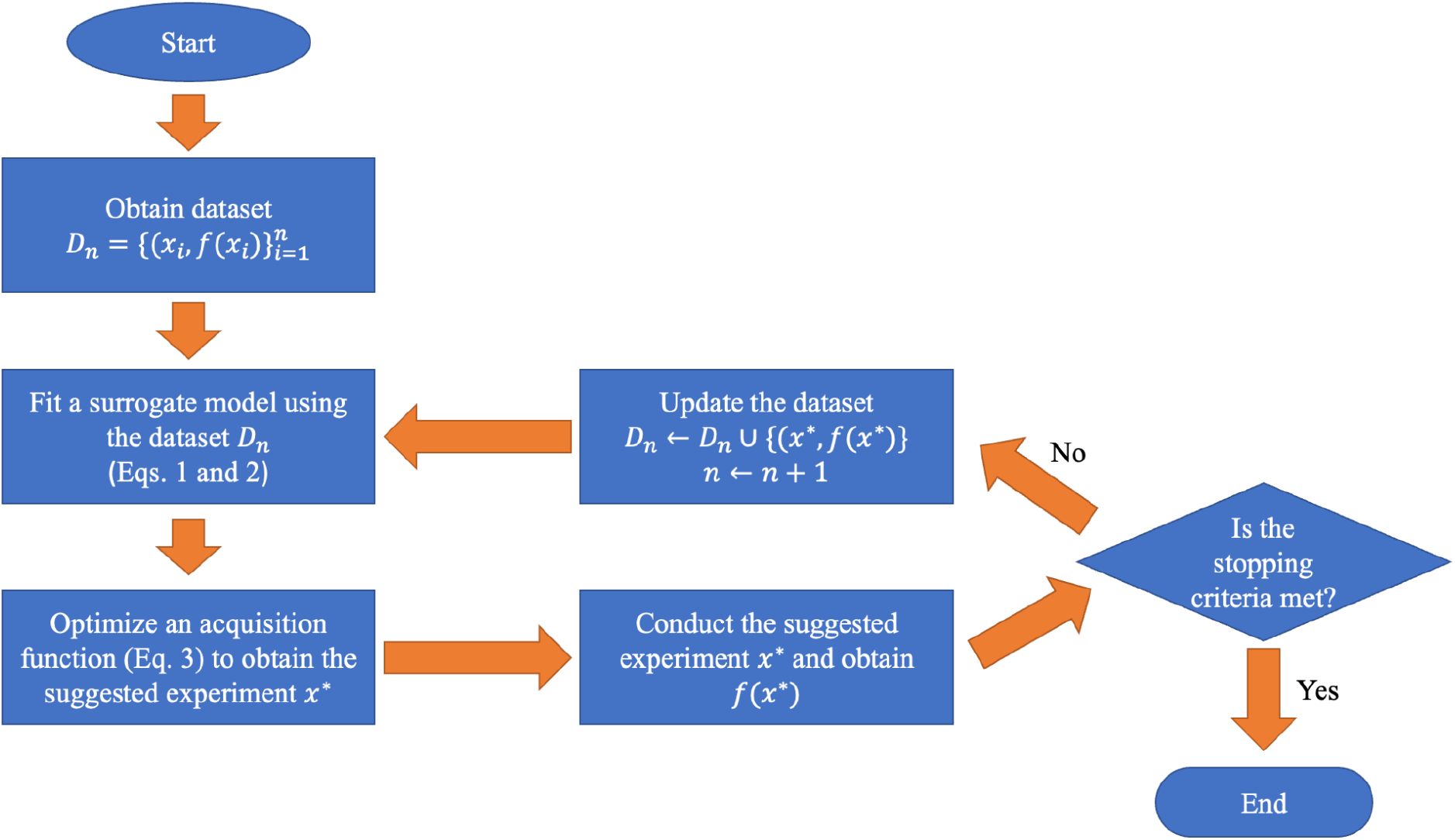
Workflow of the Bayesian Optimization algorithm

### 2.3 Optimization parameters: Complexation efficiency, yield, and loading capacity

The efficiency of forming the complex was evaluated by finding the complexation efficiency and yield percentage. A low value in either parameter indicates a loss of the initial substrate, which would result in escalated costs. To calculate these parameters, a Pierce™ Rapid Gold Bicinchoninic Acid (BCA) Protein Assay Kit was used to quantify free phytase concentration in the supernatants collected by centrifugation after complex formation, and these results were analyzed against the BSA standard curve (0-2 mg/mL). The data was collected after 5 min incubation at room temperature and measured at 480 nm in a 96-well plate reader (SpectraMax iD3, Molecular Devices, CA, US). The following equation was used to calculate complexation efficiency (**Eq.4**):

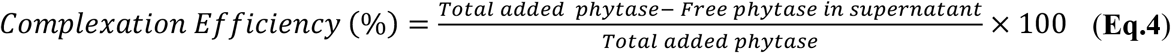

The following equation was used to calculate yield (**Eq.5**):

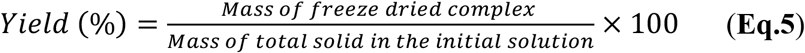

We used loading capacity to assess the efficiency of the complex by determining the amount of the target complexed or loaded in the new product. Loading capacity can be measured with the following equation (**Eq.6**):

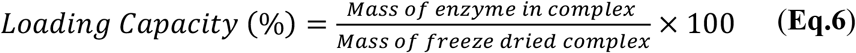

### 2.4 Phytase activity assay

Phytase activity was measured following the method outlined in the previous studies (Yang et al., 2024). The chitosan-phytase (CS-phyA) complex and control were dissolved in DI water for the heat treatment and then fully released in pH 2.0 buffer KCl-HCl (0.01 M) for gastric digestion. The final sample solution is diluted with 0.1 M sodium acetate buffer at pH 5.0 (AB) for optimal enzyme activity. The diluted phyA solution (150 µL) was then used in the activity assay.

#### 2.4.1 Thermal stability

We used a phytase activity assay to assess the phyA activity after the heat treatment. The concentration of the phyA was controlled to be the same as the CS-phyA complex to ensure an accurate comparison in thermal stability. The phyA and CS-phyA solution in DI water (~pH 7) was heated at 100 °C for 12 min in a water bath to replicate the cooking process. The residual activity percentage can be calculated using the following equation (**Eq.7**):

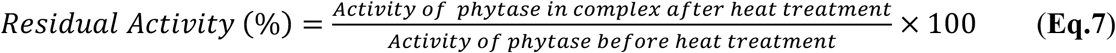

#### 2.42 Enzyme release profile

The amount of phytase released from the complex was measured at three different temperatures, 25, 50, and 100 °C, each temperature at three different time points: 20, 40, and 60 min. The released phytase was quantified using a Pierce™ Rapid Gold Bicinchoninic Acid (BCA) protein assay kit and measured as outlined in the previous study (Khongkomolsakul et al., 2025).

#### 2.4.3 In vitro digestion: Gastric stability

Using a phytase activity assay, gastric stability was assessed using Pepsin from porcine gastric tissue (2000 U/mL) in simulated gastric fluid (SGF) (KCl-HCl buffer, 0.1 M, pH 2.0) in a 1:1 v/v ratio with the CS-phyA complex solution dispersed in DI water. The solution was incubated at 37 °C for 2 h to simulate in vitro digestion (Do et al., 2016). The residual activity percentage was calculated using the following equation (**Eq.8**):

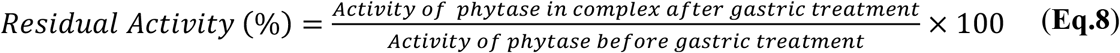

For the combination of treatments, including heat and gastric treatment, we used the same thermal stability test protocol followed by gastric digestion. The residual activity percentage was calculated as follows (**Eq.9**):

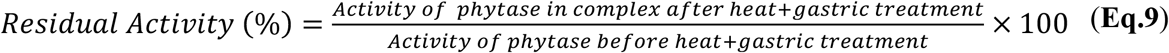

### 2.5 Complex Characterization

#### 2.5.1 Circular dichroism

Circular dichroism (CD) measurements were collected in duplicate for all samples by a JASCO-1500 spectrometer (Easton, MD, US) using a 1-cm path-length quartz cell. Thermally treated samples were pre-treated under the same conditions as the phytase activity assay. The CD spectra were recorded to confirm the protein’s secondary structure by measuring the wavelength from 190-300 nm with 1 nm resolution at each step and a 30 nm/min speed. The CD spectra were analyzed using BeStSel software (Micsonai et al., 2022).

#### 2.5.2 Confocal laser scanning microscope (CLSM)

Fluorescein 5(6)-isothiocyanate (FITC) was used as a fluorescence probe to visualize the microstructure. For protein labeling, before complexation, 50 mL of (2.5 mg/mL) phyA in pH 9.0 carbonate buffer (10 mM) was mixed with 1 mL of (2.5 mg/mL) FITC at a ratio of 50:1 at 4 °C for 48 h. The mixture of FITC-labeled phyA was filtered with a column filled with Sephadex G-25 (size exclusion chromatography resin cut off: Mw 1000 – 5000) and analyzed using UV Vis spectroscopy. The complex morphology was captured using a CLSM (i880, Carl Zeiss, Göttingen, Germany) with excitation and emission wavelengths of 488/522 nm for FITC.

#### 2.5.3 Scanning electron microscope (SEM)

To confirm the particle size and the complex formation morphology, the freeze-dried powder samples were examined using a Zeiss Gemini 500 Scanning Electron Microscope (SEM) (Jena, DE) with the same settings as the prior study (Khongkomolsakul et al., 2025).

### 2.6 Molecular docking

Molecular docking was used to observe the type of intermolecular interactions between amino acids on phytase (phyA) and chitosan (CS) to confirm the interactions at different phytase pH protonation states. The phytase structure was retrieved from the Protein Data Bank (PDB code: 3K4P) (Oakley, 2010). The structure was obtained through x-ray diffraction with a molecular weight of 99.11 kDa. To perform molecular docking for phytase, the AutoDock tool 1.5.7 software was used to eliminate water and merge non-polar hydrogen before the change in protonation state (Sanner, 1999). We then used the CHARMM-GUI and adjusted the protonation state for the system at pH 2.0 and 7.0 (http://www.charmm-gui.org/) (Jo et al., 2008; Park et al., 2023). The CS structure of 6 D-glucosamine (GlcN) units crosslinked with beta-1,4 glycosidic bonds was built and optimized using Avogadro Version 1.2.0n. The CS structure was added with Gasteiger charges, and all torsions were fixed with the AutoDock tool 1.5.7.

The molecular docking was performed using blind docking with a 126 Å cube grid at 1.000 Å spacing on AutoDock Vina 1.2.3 software (Eberhardt et al., 2021; Panda et al., 2020; Trott & Olson, 2010). Discovery Studio visualization software (BIOVIA, Dassault Systèmes, San Diego, v24.1.0) and ChimeraX were used to visualize the docking structure (Pettersen et al., 2021).

### 2.7 Statistical analysis

The data were presented as mean values with standard deviations in triplicate for all the activity and protein measurements and duplicates for circular dichroism measurements.

Statistical analysis was done using GraphPad Prism 10 and JMP Pro 16 using Tukey HSD comparison test with a significance level of p < 0.05.

## 3. Results and Discussion

### 3.1 Complex Optimization: pH

The activity recovery of all commonly used anionic polysaccharides to form a coacervation using electrostatic interaction has been studied using a combination of experimental work and molecular docking (Khongkomolsakul et al., 2025). At the optimum formation condition for phytase immobilization with anionic polysaccharides, which require a low pH level, are not able to stabilize the enzyme when heated. Therefore, the target pH for Chitosan-phyA complex formation was chosen to be at a neutral pH, which is closer to that used in food matrices. We used a pH shifting method to form the complex to ensure that the complex was stable at neutral pH and would achieve thermal protection for the enzyme for 10 min of thermal treatment at 100 °C. Parameters used to identify the optimized condition for complex formation included complexation efficiency, loading capacity, yield, and, most importantly, the enzyme’s stability under specific heat or gastric conditions.

The phyA has an isoelectric point of approximately 5.0 and exhibits optimal activity at pH levels between 2.0 (50% activity) and 5.0 (full activity) (Dotsenko et al., 2022). Consequently, in this study, we used a pH range of 2-7 to mitigate any impact on the enzyme activity. Within this pH range, phyA has a negative zeta potential, and CS is positively charged between pH 5 and 7 **(Figure 3a)**. Thus, we compared the formation of the final pH at 5-7 with 1:2, 1:1, and 2:1 ratios (CS: phyA). Complex formation was observed at pH 7 but not at pH 5 or 6. After centrifugation, a complex formation was observed by optical imaging, and under pH 7.0 conditions, a higher complex yield was obtained in comparison to pH 5 or 6 **(Figure S1)**. Therefore, the final pH of 7 was selected.

**Figure 3.**
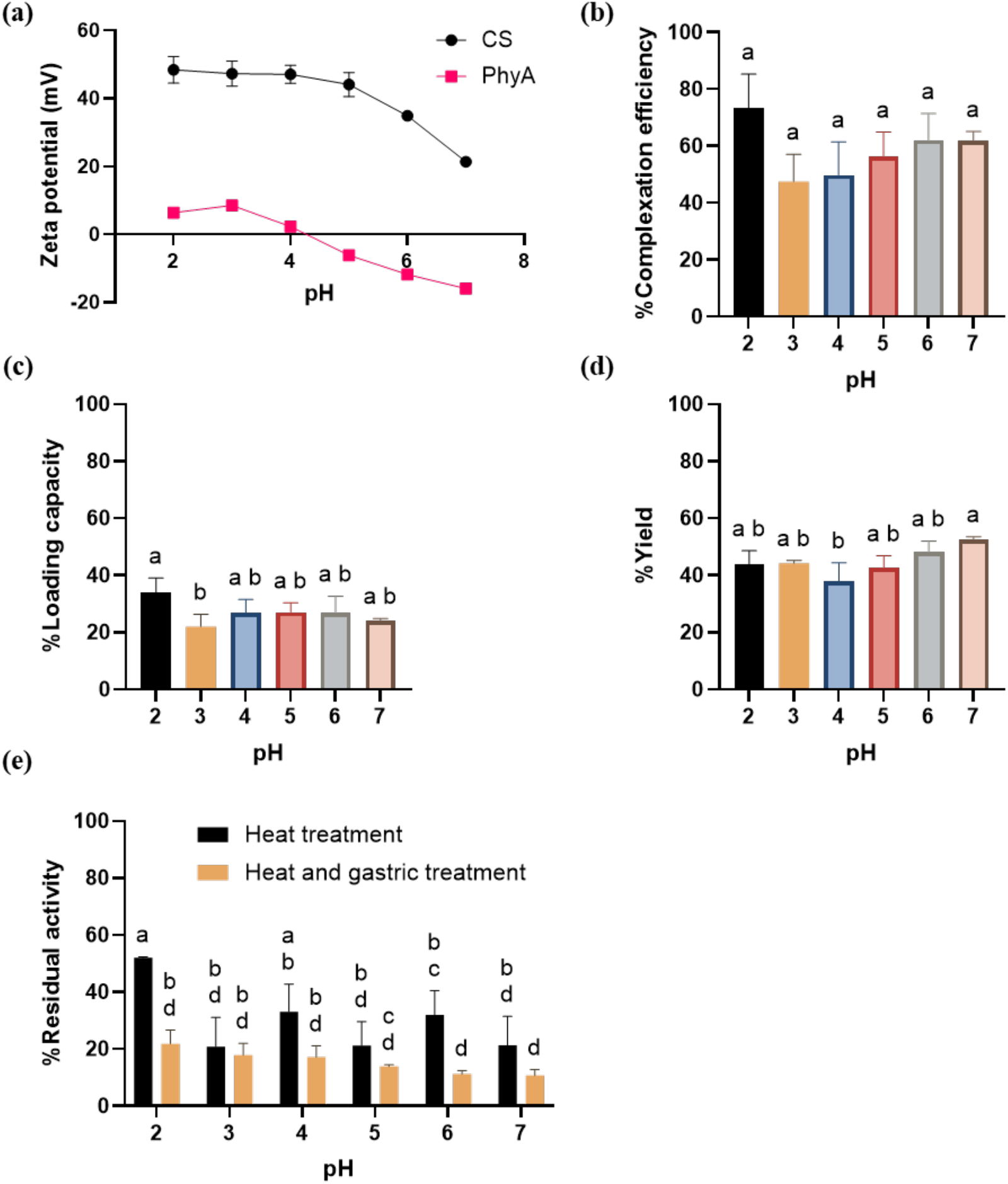
Complexation data for 1:1 CS-phyA rations measured at pH 7 after pH shifting from initial pH: (a) Zeta potential of CS and phyA, (b) complexation efficiency, (c) loading capacity, (d) complex yield, and (e) residual activity of enzyme after thermal and thermal with gastric treatment.

To determine the initial pH condition, we used a 1:1 ratio and tested a range of pH levels from 2 to 7. This initial pH condition was implemented to blend the CS and phyA solution for 30 minutes before adjusting the pH to the final level of 7. The complexes’ complexation efficiency, loading capacity, and yield formed at different starting pH levels were measured at pH 7, and for most conditions, they did not show significant differences, aside from the yield at pH 4 and 7 and the loading capacity at pH 2 and 3 (**Figure 3b-d**). At pH 2, higher complexation efficiency and loading capacity values were observed. The complexes formed at low pH levels were then shifted to pH 7 and tested for thermal stability. The complex at pH 2 exhibited significantly (p < 0.05) greater thermal stability than the complexes formed at other pH levels, except for pH 4 **(Figure 3e)**. Although not significantly higher than pH 4 (p < 0.05), the residual activity value for the complex formed at pH 2 remains the highest residual activity at ~50%. This can be attributed to the solubility of CS. In a more acidic environment, the amino group gains a hydrogen ion (H^+^) and becomes positively charged (−NH_3_^+^) due to CS’s pKa of 6.5 (D. Li et al., 2022). This alteration in pH opens up the structure of CS and provides a readily available binding site for phyA as the pH decreases. Thus, we chose pH 2 as our initial pH and pH 7 as our final pH for further optimization and characterization.

### 3.2 Complex Optimization: Ratio of chitosan to phyA

Similarly to pH optimization, we used complexation efficiency, loading, yield, and enzyme stability to select the optimized complex for further characterization. In the case of complexation efficiency, an equal ratio (1:1) of CS to phyA showed the highest complexation efficiency at 69%, with no significant difference (p < 0.05) between the 1:4, 4:1, and 8:1 CS to phyA complexes **(Figure 4a)**. CS is generally recognized as safe (GRAS) with a recommended limit of 36 mg per 60 g of baked goods and 60 mg per 60 g of plant protein products (Abourehab et al., 2022; Carlson, 2022). For food applications, therefore, the ideal complex would have a loading capacity of at least 20 w/w% for phyA for efficient use in food applications (Monnard et al., 2017). With this requirement, the 1:12-1:1 ratios of CS to phyA showed a promising loading amount of phyA in the complex with the minimum loading capacity of 28% at 1:1 **(Figure 4b)**.

**Figure 4.**
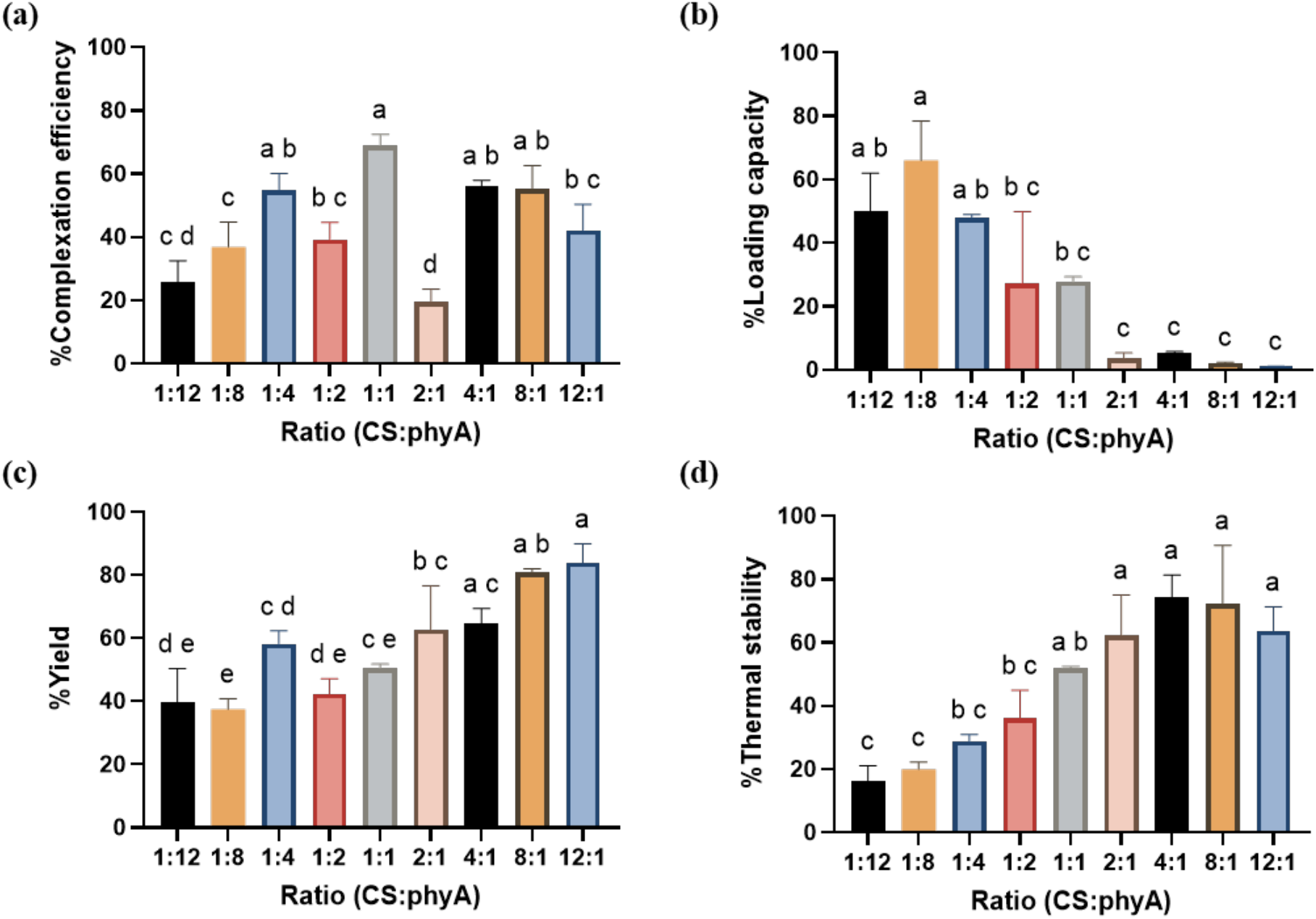
Complex at an optimized pH condition (initial pH of 2 and final pH of 7) (a) encapsulation efficiency, (b) loading capacity, (c) yield, and (d) thermal stability (residual activity after thermal treatment) of complex CS-phyA at different ratios.

The highest yield was obtained at the ratio of 12:1 CS to phyA (83% yield of complex), and at the minimum loading of 1:1 CS to phyA, the complex yield was 50% (**Figure 4c)**. The complex yield was directly related to the CS concentration. As for thermal stability, we observed that as we increased the amount of CS, the thermal stability increased and started to decrease after 4:1 CS to phyA due to repulsion between CS. The complex ratio with the highest thermal stability was 4:1 CS to phyA, as observed by the residual activity. For 4:1 CS to phyA, the residual activity was 74%, and no significant difference was found between 1:1 to 12:1 CS to phyA ratios **(Figure 4d)**. The two ratios that demonstrated the best thermal stability, exceeding 50%, were 4:1 and 1:1 CS to phyA. The 1:1 complex was deemed suitable due to its ability to meet the loading capacity requirement and maintain substantial residual activity following heat treatment. In contrast, the 4:1 ratio was selected for its exceptional thermal stability (74%) and its potential for subsequent activity assessment post-gastric digestion. To further understand the effect of the CS to phyA ratio, we further examined the release profile of the complex at various elevated temperatures with these 2 ratios..

### 3.3 Complex Optimization: Bayesian optimization (BO)

In this section, we explore the use of Bayesian Optimization (BO) for guiding experimental design in the context of the complexation efficiency of phyA. In our preliminary experiments, we evaluated complexation performance across a range of pH conditions and observed that pH 2.0 yielded the most promising results in terms of thermal stability. Based on this finding, we constrained our study to this fixed pH condition and focused on optimizing the mixing ratio of CS to phyA to maximize thermal stability.

A total of eleven mixing ratios (CS:phyA)—1:12, 1:8, 1:6, 1:4, 1:2, 1:1, 2:1, 4:1, 6:1, 8:1, and 12:1—were experimentally tested. The optimal ratio, based on our experimental results, was found to be 4:1 (CS:phyA). To assess the potential of BO in efficiently identifying this optimal condition while minimizing the number of experiments needed, we constructed a retrospective benchmarking study using the complete experimental dataset. In this setup, we simulated a scenario in which the optimal ratio is not known a priori and applied BO to sequentially select ratios for evaluation.

To run the BO, we encoded the ratios by the amount of CS per unit phyA and formed the search space *χ* = {0.08, 0.13, 0.17, 0.25, 0.50, 1.00, 2.00, 4.00, 6.00, 8.00,12.00}.The BO algorithm is initialized with two randomly selected mixing ratios and uses the GP model and the EI acquisition function to guide the selection of subsequent candidates. The procedure continues until we until all considered ratios have been experimentally tested. To account for potential bias introduced by initialization, we evaluate BO across all 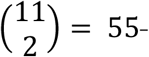 possible combinations of initial ratio pairs and report average performance.

As a baseline, we include a random selection strategy (Random), which begins with the same initial pairs as BO but selects subsequent ratios uniformly at random, without using an acquisition function, and uses the same stopping criteria.

The optimization progress curves show the best thermal stability observed versus the number of experiments conducted **(Figure 5)**. The results indicate that for a fixed number of conducted experiments, BO consistently identifies a better CS to phyA ratio that yields higher thermal stability, than the Random baseline. We also calculated the average number of experiments required for BO and Random baseline to identify the optimal ratio 4:1 (CS:phyA) or equivalently input x = 4.00 over all 55 trials and found that BO requires, on average, only 4 experiments compared to 7 required by Random. These findings highlight the value of BO as a practical strategy for experimental optimization, offering a better experimental condition while requiring a smaller number of experiments.

**Figure 5.**
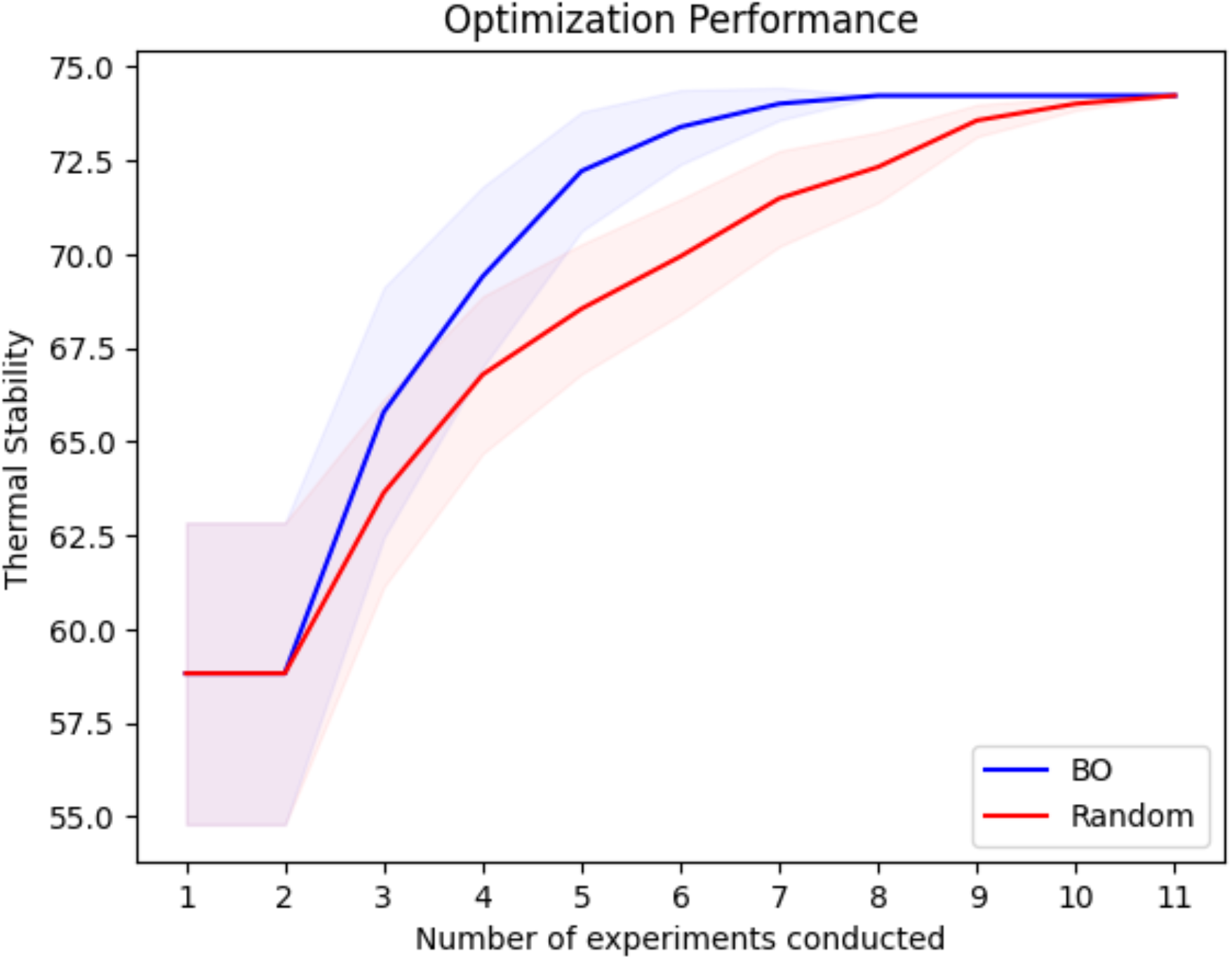
Optimization progress curves comparing BO and Random algorithms to find the optimal ratio for pH 2.0. Solid lines represent the mean across 55 trials, and shaded regions denote standard errors

### 3.2 Effect of complexation on enzyme stability

#### 3.2.1 Effect of thermal treatment

The release profile was used to evaluate the improvement in thermal stability of phyA when complexed with CS. Our study involved measuring the release profile of the complex at 4:1 and 1:1 CS to phyA ratios at 25, 50, and 100 °C over 60 min, with the amount of protein release readings through BCA assay taken at 20-min intervals (**Figure 6a and 6b, respectively**). At the 20-min mark, the 4:1 CS to phyA complex released less than 20 w/w% at all temperatures, while the 1:1 CS to phyA complex exhibited a slightly faster release rate at 100 °C, with 41 w/w% released in 20 min. The release profile correlated with the residual activity of the complex following heat treatment at 100 °C (**Figure 6c**). As the complex was incubated at a higher temperature, the enzyme was trapped in the system, which prevented it from unfolding, and the phytase structure remained unchanged. This resulted in a high retention of enzyme activity as the secondary structure remained intact and the active site was retained and available for binding with the substrate.

**Figure 6.**
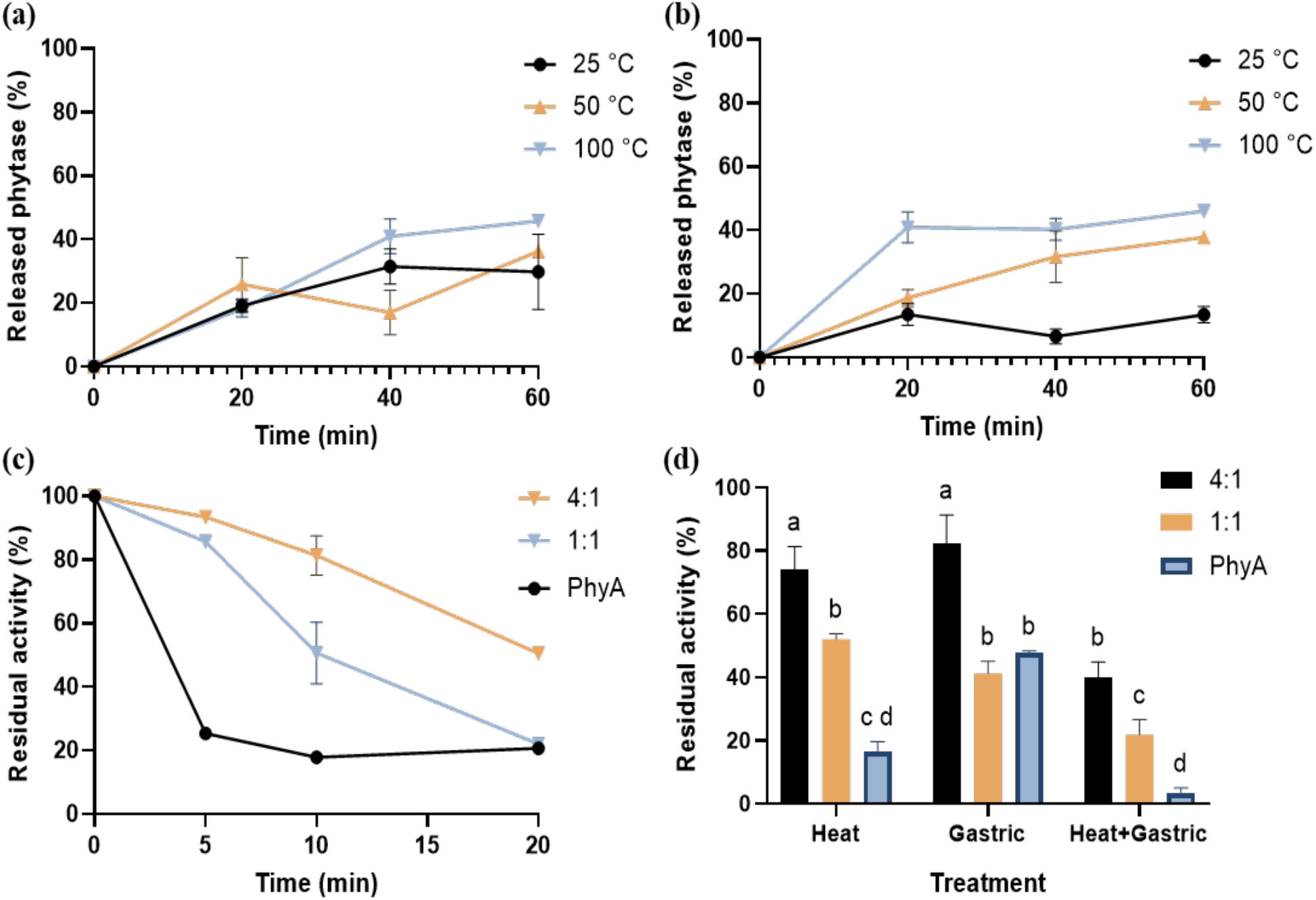
Releasing profile of complex CS-phyA (a) 4:1 and (b) 1:1 at 25, 50, and 100 °C at different time points in DI water, (c) residual activity after thermal treatment at different time points, and (d) residual activity with thermal treatment at 100 °C for 12 minutes, gastric treatment, and the combined treatment for phyA (control), and complexes of 4:1 and 1:1.

During the initial 5 min, the control sample (phyA without CS) demonstrated a significant decline (p < 0.05), retaining only about 20% of its original activity after heat treatment. In contrast, the complex samples exhibited higher enzyme activity retention rates, with the 4:1 and 1:1 CS to phyA complexes retaining approximately 90% and 85% of their residual activities, respectively. The trend continued to the 10-min mark, where the phyA control and the 1:1 and 4:1 CS to phyA complexes retained their residual activity levels of 18%, 51%, and 81%, respectively. Finally, after 20 min of incubation at 100 °C, the residual activity was about 22% for both phyA and 1:1 CS to phyA, while the 4:1 CS to phyA complex retained a higher residual activity level of 51%. These findings suggest that CS complexation of phyA protected phyA from heat treatment.

#### 3.2.1 In vitro digestion

The phyA control sample showed 3% residual activity after a combined thermal and gastric treatment **(Figure 6d)**. Comparatively, after a combined thermal and gastric treatment, the CS phyA complexes retained 40% residual activity when the CS to phyA ratio was 4:1, and the 1:1 CS to phyA complexes showed 22% residual activity. This represents a 13-fold increase in residual activity compared to the enzyme itself. The decrease in residual activity by about half between the 4:1 and 1:1 CS to phyA complexes correlated with the release profile, as the acidic conditions enhanced CS solubility, allowing pepsin to partially denature the released phyA in the gastric simulation.

### 3.3 Characterization of optimized complex

#### 3.3.1 Circular dichroism

Circular dichroism (CD) allows for the evaluation of protein secondary structures, which helps to identify protein folding and stability. A positive band at 193 nm and negative bands at 208 and 222 nm are major indicators for proteins with mainly α-helical structure (Greenfield, 2006). We measured the CD spectrum for the enzyme itself and the complex of 4:1 and 1:1 CS to phyA, both before and after thermal treatment. Phytase from *A. niger* has a similar structure as phyA, and the structure as obtained from PDB (3K4P) showed a secondary structure comprising approximately 40% α-helices, 30% β-sheets, and 30% other structures. A strong negative band at 222 nm is indicative of phyA’s α-helix structure (**Figure 7a**). Before the thermal treatment, phyA had 59% α-helices and 41% of β-sheets; after thermal treatment, the ratio of α-helix decreased to 42% with an increase of β-sheets to 58%. This reduction in the α-helix to β-sheet ratio of phyA was associated with decreased enzyme activity after the thermal treatment and was previously reported by other studies (Correia et al., 2008).

**Figure 7.**
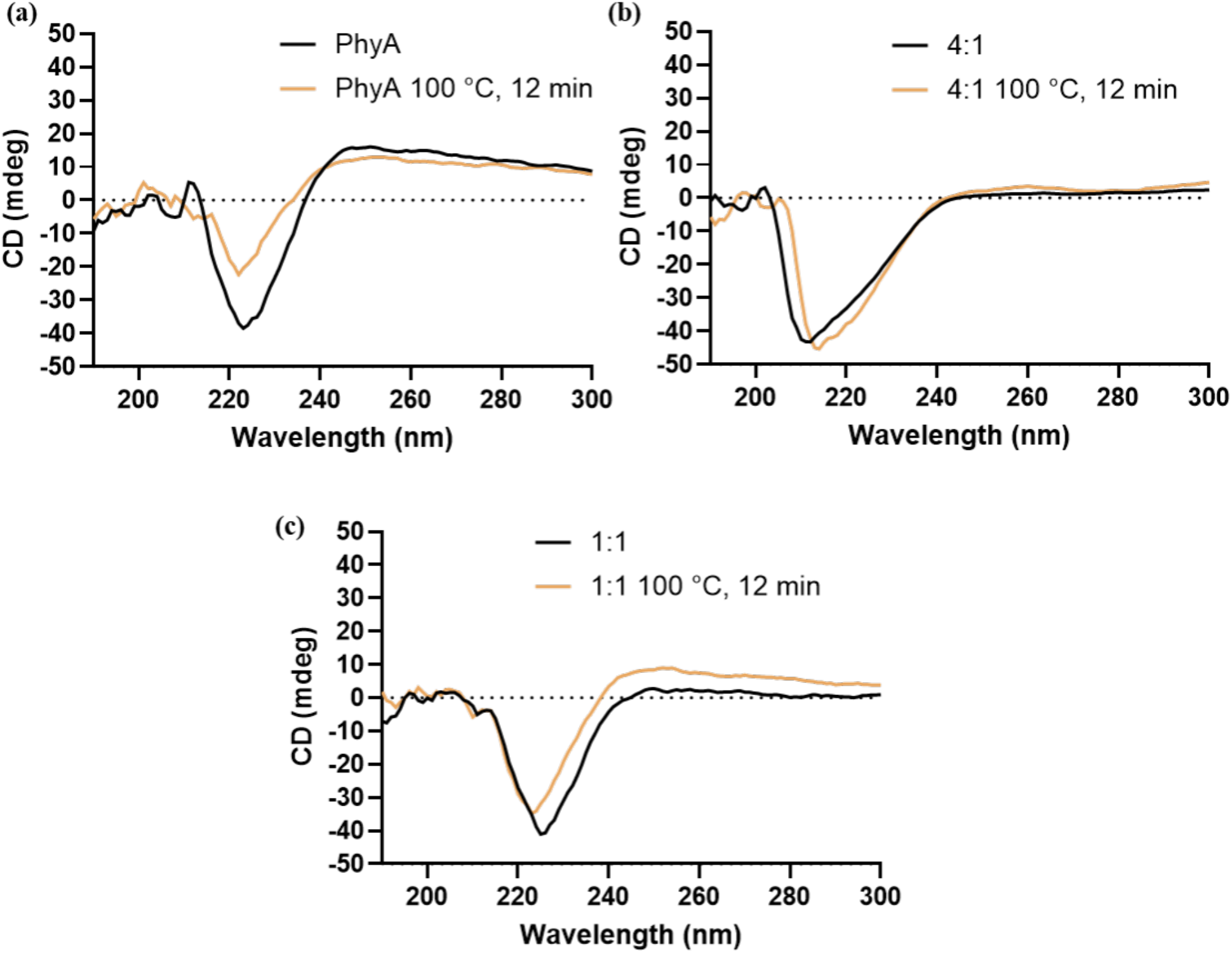
Circular dichroism graph of (a) phyA, complex (b) 4:1, and (c) 1:1 before and after thermal treatment.

Meanwhile, the 4:1 CS to phyA complex showed 74% α-helices and 26% β-sheets before thermal treatment, and after thermal treatment, the ratio of α-helix was 62% and with a slight increase to 38% for β-sheets **(Figure 7b)**. The 1:1 CS to phyA complex had 60% of α-helix and 40% of β-sheet before thermal treatment, and remained unchanged after thermal treatment **(Figure 7c)**. The CD spectra of the complexes in both 4:1 and 1:1 CS to phyA ratio when compared to phyA after treatment, did not show a similar decrease in α-helices indicated by the band at 222 nm, this indicates that the complexes were more stable with respect to the thermal treatment (Yang et al., 2024). This stability in the phyA’s secondary structure in the complex after thermal treatment, combined with the release profile, verified the complex’s ability to protect the enzyme and retain enzyme activity upon heat stress.

#### 3.3.2 Morphology

The SEM images reveal the CS, phyA, and the 4:1 complex structure at 10 and 2 µm, respectively **(Figure 8a-f)**. CS exhibited a fibrous-like structure compared to phytase, which has a smooth surface in a powder form, but after complexation with phyA, particle agglomeration occurred on the surface. This could be attributed to pH-induced self-aggregation of CS, facilitating the complexation of phyA within the structure.

**Figure 8.**
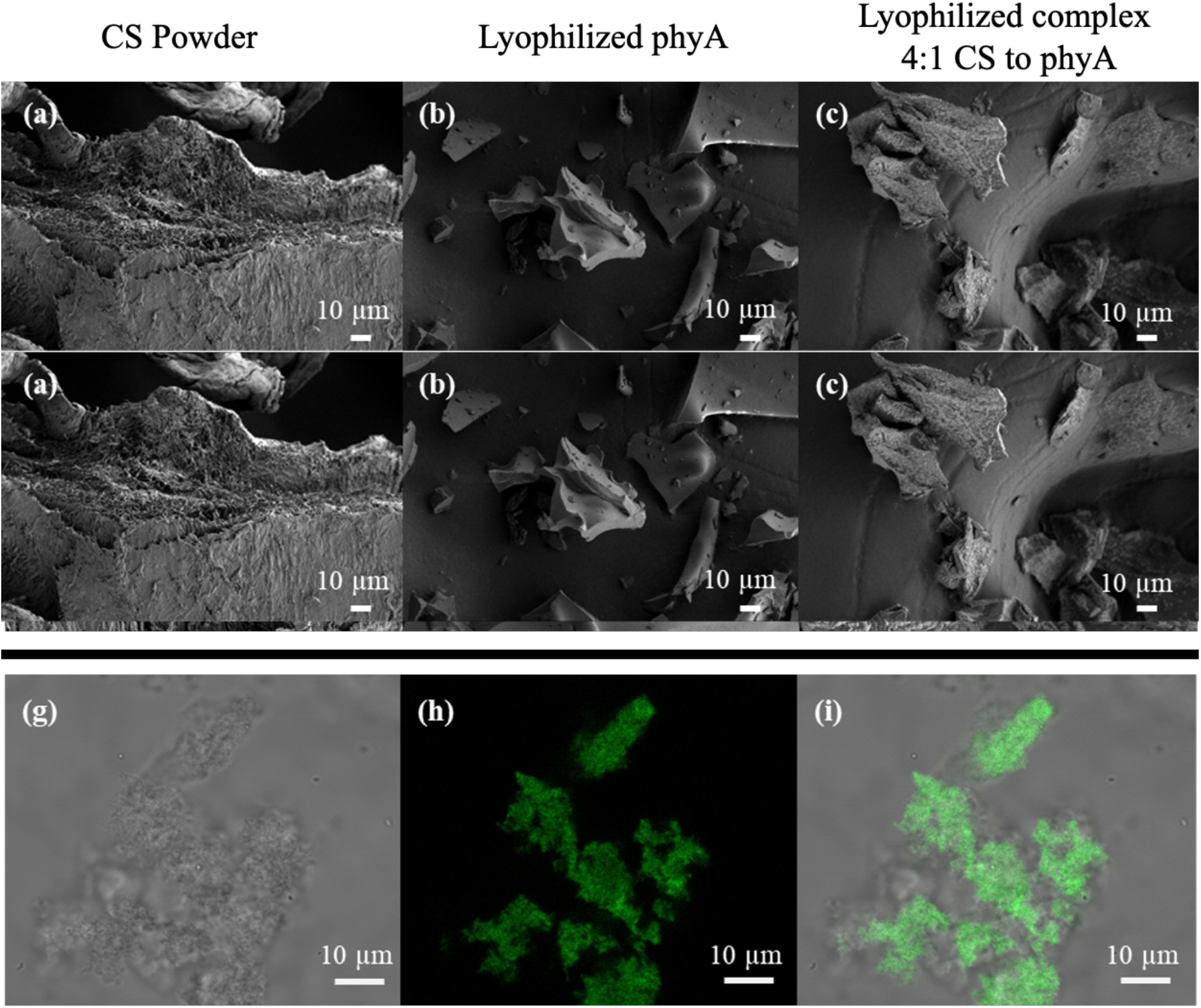
SEM images of (a, d) CS powder, (b, e) lyophilized phyA, (c, f) lyophilized complex 4:1 CS to phyA at 10 and 2 µm, respectively, and (g, h, i) CLSM images of complex 4:1 CS to phyA.

CLSM images of the 4:1 CS to phyA complex **(Figure 8g-i)** confirmed the presence of labeled protein inside the CS particle (Weng et al., 2022). PhyA was labeled with FITC, and excess FITC was removed using a Sephadex G-25 column prior to complexation. Additionally, Z-axis imaging showed that the FITC-labeled phyA was more intense in the middle of the complex as we sliced through the z-axis **(Figure S2)**. This provided conclusive evidence that phyA was indeed located at the center of the CS microparticles.

### 3.4 Molecular docking

Molecular docking was used to observe the best conformation of CS, and 3K4P was used as a representative of phytase from *A. niger* (phyA) in different protonation states of phyA **(Figure 9)**. Blind docking was used and performed using Autodock Vina, using the scoring function to determine the best conformation. Docking of phyA at a protonation state of pH 7 was performed, and the binding site was found to be close to the active site of phyA **(Figure 9a, b)**. In our previous study, we saw a lower activity recovery for the coacervate complex with similar best conformation from blind docking (Khongkomolsakul et al., 2025). The result was similar in the CS microparticles docking with phyA at pH 7 because we did not see the enzyme’s activity, as it takes time to release in DI water at room temperature **(Figure 6a)**. That means, in the context of activity recovery, we also had to consider the release profile carefully to match the application. We can see that after the protonation state changed from pH 7 to pH 2 **(Figure 9c, d)**, the interactions decreased, as well as the best conformation changed. Thus, we saw a high activity retention rate in phyA after the thermal treatment. This knowledge of polysaccharide-enzyme molecular docking may open up a novel approach to selecting the type of carrier and how to predict the protection and enzyme activity after the complex formation.

**Figure 9.**
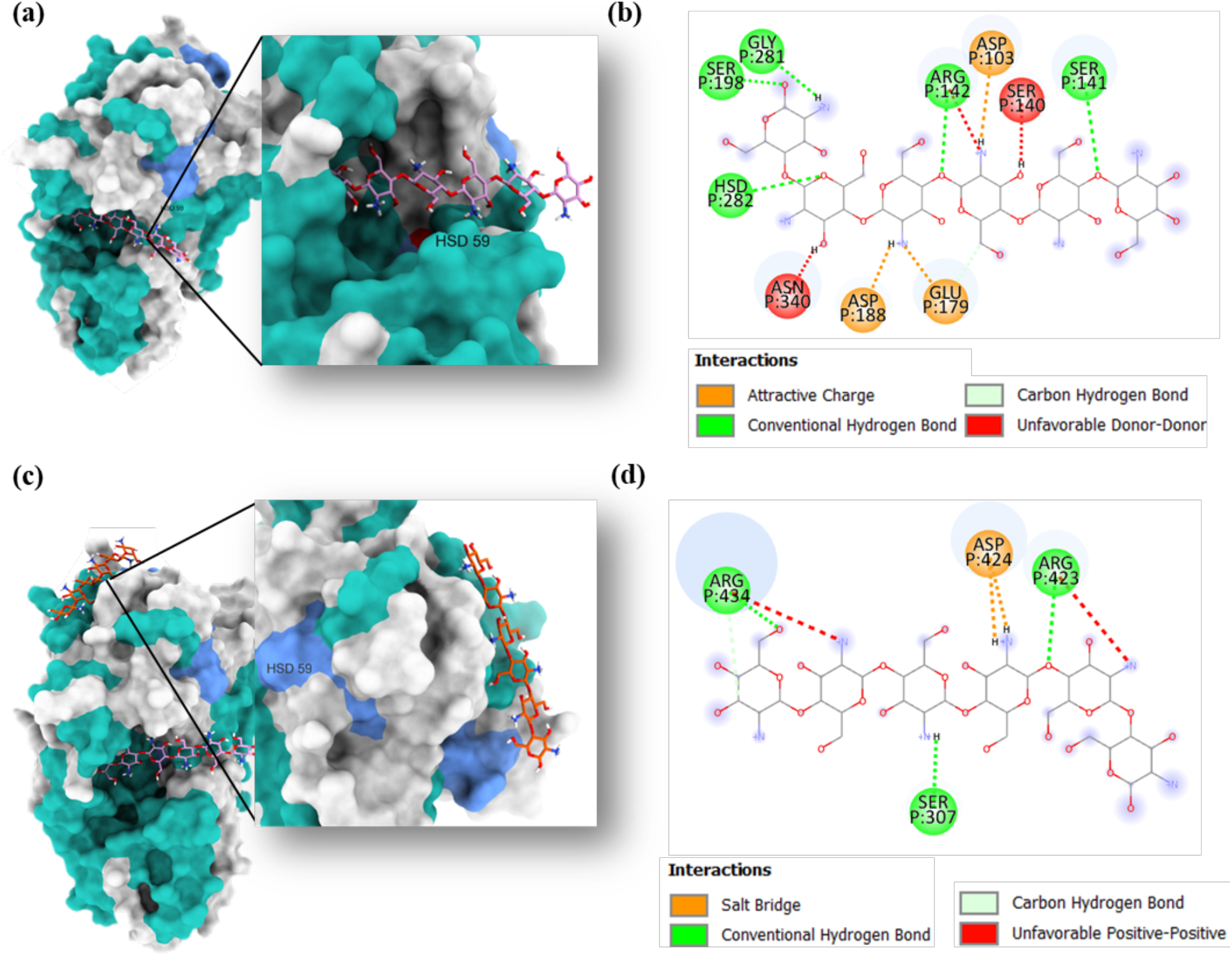
(a) Molecular docking structure of phyA (3K4P) at pH 7 protonation state and CS (labeled in pink) with the best binding affinity at −47.2 kJ/mol, (b) intermolecular interaction between phyA (3K4P) at pH 7 protonation state and CS, (c) Molecular docking structure of phyA (3K4P) at pH 2 protonation state and CS (labeled in orange) with the best binding affinity at −46.6 kJ/mol, (d) intermolecular interaction between phyA (3K4P) at pH 2 protonation state and CS. The secondary structure of phytase was labeled by colors, in which grey dictated the coils, light sea green dictated the helixes, and cornflower blue dictated the strands. The intermolecular interactions were also labeled in color, in which green colors represented hydrogen bonds, including Van der Waals and C-H bonds, orange represented an attractive charge, and red represented an unfavorable interaction, including charge repulsion and acceptor/donor clash.

## 4. Conclusions

Stabilizing phyA at practical use temperatures up to 100 °C has been difficult, and there is limited research on residual activity at different time intervals, potentially compromising thermal stability results. This study introduces an innovative method for complexing enzymes, specifically phytase (phyA), within micro-sized particles of chitosan (CS). Using a pH shifting approach, we were able to find a suitable starting pH for maximum yield and loading capacity and then shift the pH to food-appropriate levels while protecting phyA within chitosan microparticles. Bayesian optimization (BO) was able to reach the best ratio of CS to phyA in fewer iterations and fewer suggested experiments than random or grid search. These promising results highlight the potential of using BO for general parameter optimization or optimizing intermediate parameters for any complexation system (Buathong et al., 2024). This study paves the way for future food applications to use BO and to expedite optimization processes.

The optimized complex initially formed at pH 2 and shifted to pH 7, and a mass ratio of 4:1 CS to phyA showed significant (p < 0.05) enhancement in thermal treatment, retaining up to 74% residual activity after complexation, and then 40% residual activity after heat and gastric treatment. This substantial improvement will enable the use of phyA in high-phytate food matrices, thereby improving micronutrient absorption for human consumption. For future research, enzyme activity will be further enhanced through ternary complexation, and an alternative formation strategy for scaling up will be explored. This allows for cost-effective production through comparative assessments of activity retention and cost.

## Data availability statement

Data associated with the figures in this manuscript can be found at:

https://doi.org/10.5281/zenodo.15243436. Supporting information contains additional figures.

## Declaration of competing interest

The authors have no conflict of interest to declare.

## Author credit statement

**Waritsara Khongkomolsakul:** Conceptualization, investigation, data acquisition, formal analysis, visualization, writing original draft, reviewing, and editing. **Poompol Buathong:** Bayesian optimization, writing original draft, reviewing, and editing. **Eunhye Yang:** Conceptualization, writing, reviewing, and editing. **Younas Dadmohammadi:** Supervision, funding acquisition, reviewing, and editing. **Yufeng Zhou:** Data acquisition (Circular dichroism). **Peilong Li:** Data acquisition (CLSM). **Lixin Yang:** Data acquisition (particle distribution). **Peter I. Frazier:** Bayesian optimization, writing, reviewing, supervision, and editing. **Alireza Abbaspourrad:** Project conceptualization and administration, supervision, funding acquisition, writing, reviewing, and editing

## Acknowledgments

This research received support from the Nutrition Research and Development Program (INV-043930) sponsored by The Bill & Melinda Gates Foundation in Washington, US. The electron microscopy facility at the Cornell Center for Materials Research (CCMR) was utilized in this study, which was funded by the National Science Foundation Materials Research Science and Engineering Centers (MRSEC) program (DMR-1719875). The CLSM imaging data was acquired through the Cornell University Biotechnology Resource Center (RRID: SCR_021741) with NIH S10RR025502 funding. The authors also acknowledge Kelley J. Donaghy for her assistance editing this manuscript.

## Notes

https://doi.org/10.5281/zenodo.15243436

### Competing Interest Statement

The authors have declared no competing interest.

