## Supplemental Figures for "Improving Thermal and Gastric Stability of Phytase via pH Shifting and Coacervation: A Demonstration of Bayesian Optimization for Rapid Process Tuning"

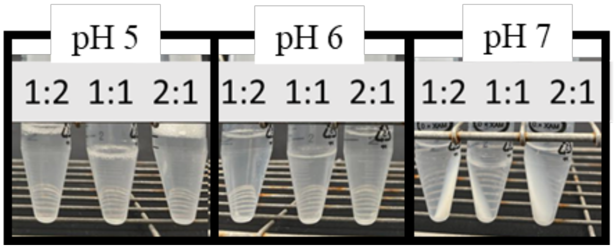


**Figure S1.** The optical image of the centrifuged complex at different final pH (5-7) at 1:2, 1:1, and 2:1 ratio.

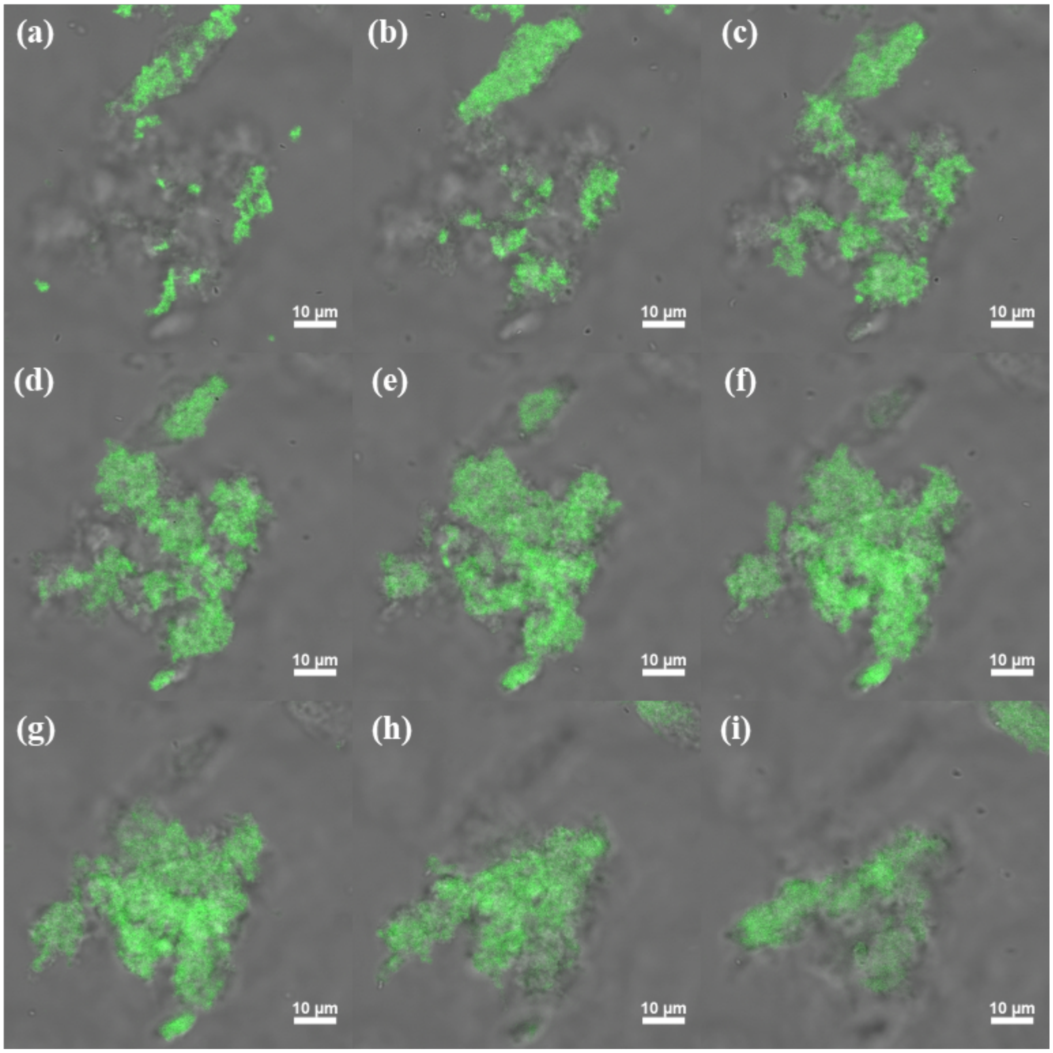


**Figure S2.** CLSM representative images of complex C-phyA 4:1 at 10 μm in different Z-axis cross-section slices from top to bottom (a-i). The video of all the CLSM stacked images (42 images) will be available in the supplementary file.
